# Gell: A GPU-powered 3D hybrid simulator for large-scale multicellular system

**DOI:** 10.1101/2022.09.01.506296

**Authors:** Jiayi Du, Yu Zhou, Lihua Jin, Ke Sheng

## Abstract

As a powerful but computationally intensive method, hybrid computational models study the dynamics of multicellular systems by evolving discrete cells in reacting and diffusing extracellular microenvironments. As the scale and complexity of studied biological systems continuously increase, the exploding computational cost starts to limit large-scale cell-based simulations. To facilitate the large-scale hybrid computational simulation and make it feasible on easily accessible computational devices, we develop a fast and memory-efficient open-source GPU-based hybrid computational modeling platform Gell (GPU Cell), for large-scale system modeling. We fully parallelize the simulations on GPU for high computational efficiency and propose a novel voxel sorting method to further accelerate the modeling of massive cell-cell mechanical interaction with negligible additional memory footprint. As a result, Gell efficiently handles simulations involving tens of millions of cells on a personal computer. We compare the performance of Gell with a state-of-the-art paralleled CPU-based simulator on a hanging droplet spheroid growth task and further demonstrate Gell with a ductal carcinoma *in situ* (DCIS) simulation. Gell affords ~150X acceleration over the paralleled CPU method with one-tenth of the memory requirement.

**Author Summary:** Numerical cell simulations provide indispensable insight into the cell-to-tumor tissue transition and help reduce biological experimental variables. However, the availability and practicality of large-scale cell simulation tools have been limited by high computational cost, slow performance, or proprietary. Recent developments in open-source simulation codes and GPU implementation have partially addressed the challenge. We further optimized the cell simulation platform for GPU implementation in this work. As a result, benchmark cell simulation experiments can be performed efficiently on a personal computer with a modern GPU. We made the platform open source to encourage community adoption and collective development.

## Introduction

Computational modeling has become an important tool for studying the dynamics of tissue development and tumor response to different therapeutic interventions over the past three decades. Three major types of models are commonly utilized in these studies: discrete, continuum, and hybrid models(1). Discrete models, also known as cell-based models or agent-based models, simulate the individual behaviors and the mutual interactions of the cells in a system. Continuum models consider biological tissues as domains composed of different solid and fluid phases and describe the system evolution using partial differential equations. Hybrid models combine the aforementioned two methods as they model discrete cells in a continuum environment.

Cell-based and hybrid simulations adopt the discrete representation of the cell of interest. Compared to the continuum representation, the discrete modeling of individual cells better captures the heterogeneity of tumorous tissue with independently tracked cell states and enables a more straightforward translation from biological hypothesis to simulation rules(2). Cell-based/hybrid simulation has been utilized to explore various kinds of oncological topics(3), such as epithelial ducts and cysts through the epithelial acini models(4) (5), the development of the initial phases of avascular tumor development through multicellular tumor spheroid (MCS) models(6), angiogenesis, vascular network formation problems(7), and anti-cancer treatments effectiveness modeling(8) (9). The method for discrete representation of cells can either be lattice-based or off-lattice. Lattice-based methods are mesh-based, including cellular automaton (CA) models(10), lattice gas CA (LGCA) models(11), and Cellular Potts models (CPM)(12). Lattice-based models are susceptible to grid biases(13), which do not affect the off-lattice methods. Two major types of off-lattice models are Boundary-tracking models(14) and Center-Based models (CBMs)(1) (15). Boundary-tracking models dedicate more computational resources to the cells’ morphological dimension and are thus more computationally expensive than CBMs(2). CBMs assume a spherical cell shape and represent cell movement by displacing the cell center position. Cell movements can be realistically modeled by incorporating forces such as adhesive, repulsive, locomotive, and drag-like forces(15). Center-based representation is often a superior choice for large-scale cell-based/hybrid simulations interested in the heterogeneous development of biological tissue due to its realistic modeling of multicellular interactions and lower computational cost than other cell-based methods.

However, even with a center-based representation of discrete cells, as the scale and complexity of the studied biological systems continuously increase, the exploding computational cost still limits the large-scale cell-based/hybrid simulations. Several CPU-based cell-based/hybrid simulation software frameworks have been proposed to enable large-scale biological system modeling on high-performance computers. Biocellion(16), a closed-source commercial software, has simulated millions to billions of cells on cluster computers. Physicell(15) is an open-source parallel simulation platform capable of simulating 18.2 days of hanging drop spheroid growth with up to one million cells in 3 days on a high-performance computer. BiodynaMo(17) is another open-source parallel simulator shown 945X faster on the ‘epidemiology (medium-scale)’ benchmark using 72 CPUs on a server compared to a single thread version. Because of the demonstrated potential to accelerate cell-based simulation with parallelization, recent research has shifted to the graphic processing unit (GPU), which has an intrinsic parallel architecture with thousands of computational cores. Ya||a(18) is a paralleled agent-based model runs on GPU. Its extended spheroid cell model with spin-like polarities can simulate epithelial sheets and tissue polarity. Although Ya||a(18) can achieve 10X acceleration compared to CPU-based cell-based simulation library Chaste(19), their simulation software is not designed for large systems. Simulation of large systems using Ya||a(18) is limited by its computational complexity of O(N^2^) for cell-cell interaction. CBMOS(20) is another GPU-based software that provides a platform to study the effects of force functions, ODE solvers, time step sizes, and cellular events in CBMs. CBMOS(20) utilizes fast GPU vector operations provided by CuPy for efficient calculations and achieved a simulation speed 30X faster than their CPU version. Their emphasis, however, is on a better user interface for fast prototyping of new models. Its ability to handle large systems is still limited by the platform design, e.g., the force calculation time complexity is O(N^2^) and the GPU memory consumption can exceed 16 GB (e.g., NVIDIA Tesla T4) for >10^4^ cells. GPU BioDynaMo(21) upgrades BioDynaMo(22) by enabling GPU co-processing. With a GTX 1080Ti GPU, GPU BioDynaMo can be 130X faster than the single thread CPU only version BioDynaMo. However, GPU BioDynaMo does not solve PDEs on GPU, and its uniform grid method for force calculation still needs CPU for linked list maintenance, which significantly limits its real-world performance.

Therefore, although acceleration of cell simulation using GPU has been demonstrated, the potential has not been fully realized. Specifically, the cell-cell interaction has not been fully parallelized, and the slow data transfer between CPU and GPU results in significant overhead. In this study, we develop a new open-source fast and memory-efficient fully GPU-based hybrid simulation software, GPU cell (Gell), to overcome these bottlenecks for large-scale hybrid cell simulation.

## Methods

### Cell Model

#### Cell cycle and death

We model five cell phases in Gell. The premitotic, postmitotic, and quiescent phases are for living cells, and the necrotic and apoptotic phases are for cell death. The phase transition from premitotic to postmitotic and from postmitotic to quiescent are deterministic with a fixed gap time, respectively. Meanwhile, the phase transition from the quiescent phase to the premitotic phase and any phase transition from the living cell phase to the dead cell phase are all stochastic with a certain transition rate.

**Table 1.**
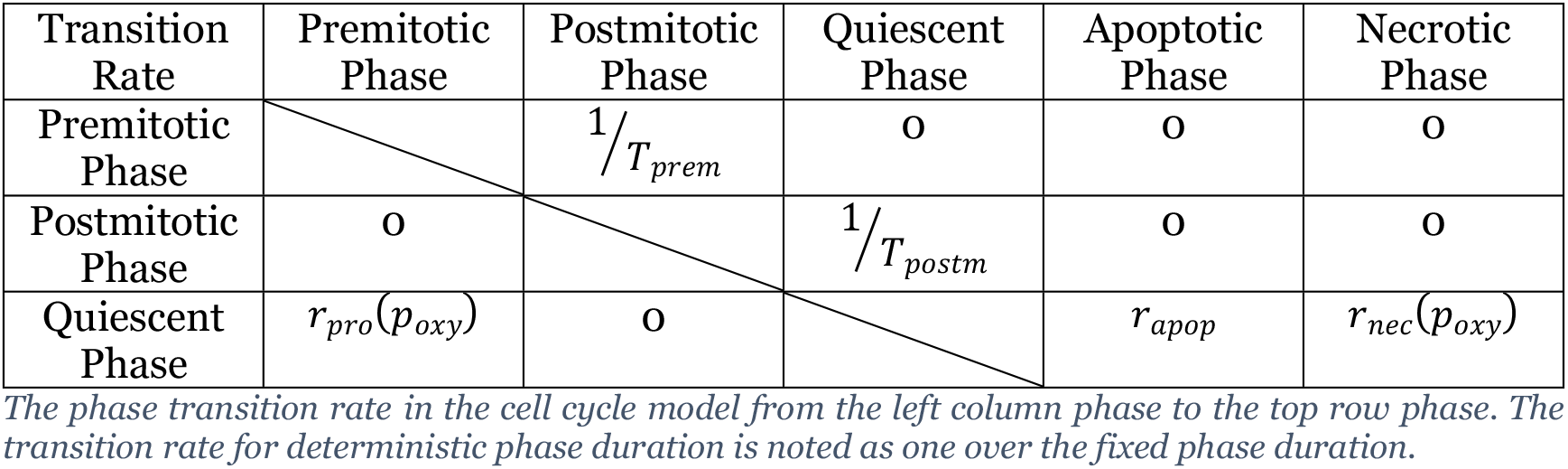
Phase transition in Gell.

The probability for any stochastic transition α to take place in a short time interval Δt is given by:

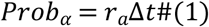

During the quiescent phase, cells maintain their standard volume and remain to be stochastically activated for division preparation at the rate *r_pro_*(*P_oxy_*):

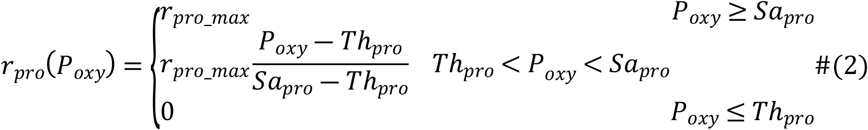

The proliferation rate increases linearly with local oxygen concentration in the given range *Th_pro_* < *P_oxy_* < *Sa_pro_*, where Th_pro_ is the minimum oxygen partial pressure required for proliferation, Sapro is the saturation oxygen partial pressure when the transition rate for proliferation reaches the maximum value r_pro_max_.

Once activated for proliferation, cells enter the premitotic phase and prepare for division. Premitotic cells gradually gain mass/volume, and then divide at the end of this phase and enter the postmitotic phase. The division process is mechanically modeled to finish in an instant, and the two daughter cells equally inherit half of the parent cell volume and be placed around the parent cell center with the displacement x_disp_(15).

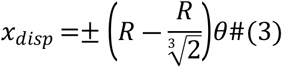

Where R is the cell radius of the parent cell, and θ is a three-dimensional random unit vector. The postmitotic phase accounts for the duration required for daughter cells to reach mechanical equilibrium and grow to be proliferation ready.

Cell death is activated stochastically for all living cells. The transition rate to enter the apoptotic phase is a constant for all cells, *r_apop_*, while the transition rate for necrotic death depends on the local oxygen partial pressure P_oxy_:

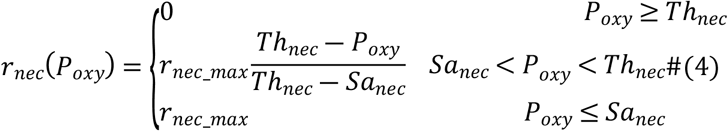

Necrosis happens only when local oxygen partial pressure is lower than the necrosis threshold Th_nec_. The transition rate increases linearly as the oxygen concentration decreases till the maximum necrosis rate r_nec_max_ is reached when oxygen partial pressure equals the necrosis saturation threshold Sa_nec_.

All cell components start to shrink for apoptotic death upon entering the phase. While for the necrotic phase, early necrotic cells first absorb fluid and swell in the oncosis process, and then enter the late necrosis process after the membrane ruptures and start to lose fluid components. Modeling of two-stage necrotic death can be critical when studying the microstructures in the hanging drop spheroid necrotic center(15). Phase transition-related parameters can be found in Tab. 2, where the listed values are all adopted from the “Ki67 Advanced” model from PhysiCell(15).

**Table 2.** Phase transition-related parameters.

| Parameter | Biophysical meaning | Reference Value |
| --- | --- | --- |
| $T_{prem}$ | Duration of premitotic phase | 13 hour |
| $T_{postm}$ | Duration of postmitotic phase | 2.5 hour |
| $r_{apop}$ | Apoptosis rate | $0.0060 \text{ hour}^{-1}$ |
| $r_{pro\_max}$ | Max proliferation rate | $0.1176 \text{ hour}^{-1}$ |
| $Sa_{pro}$ | The oxygen level when the proliferation rate reaches maximum | 10 mmHg |
| $Th_{pro}$ | The oxygen level when the proliferation rate drops to zero | 5 mmHg |
| $r_{nec\_max}$ | Max necrosis rate | 0.1667 hour<sup>-1</sup> |
| $Sa_{nec}$ | The oxygen level when the necrosis rate reaches maximum | 2.5 mmHg |
| $Th_{nec}$ | The oxygen level when the necrosis rate drops to zero | 5 mmHg |

Following Physicell(15), we divide the total cell volume V into the fluid volume V_F_ and the solid biomass volume V_S_. The solid biomass volume is further divided into total nuclear volume V_NS_ and cytoplasmatic volume V_CS_. Different cell components have different rates of volume gain and loss in different phases. All the volume changes of different cell components are modeled using ordinary differential equations (ODEs):

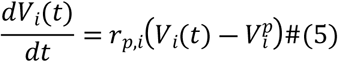

Where V_i_ is the volume of component i, r_p,i_ is the volume change rate of component i in phase p, and V_i_^p^ is the desired volume of component i in phase p. Specially, the desired fluid volume of a cell is a function of the current total cell volume V. Related parameters can be found in Tab. 3, adopted from MCF-10A human breast cancer cell line(15).

**Table 3.** Cell volume growth related parameters.

|  | Premitotic Phase | Postmitotic Phase/<br>Quiescent Phase | Apoptotic Phase | Early Necrotic Phase | Late Necrotic Phase |
| --- | --- | --- | --- | --- | --- |
| $V_F$ | 0.7502V | 0.7502V | 0 um<sup>3</sup> | V | 0 um<sup>3</sup> |
| $r_F$ | 3 hour<sup>-1</sup> | 3 hour<sup>-1</sup> | 3 hour<sup>-1</sup> | 0.67 hour<sup>-1</sup> | 0.05 hour<sup>-1</sup> |
| $V_{NS}$ | 270 um<sup>3</sup> | 135 um<sup>3</sup> | 0 um<sup>3</sup> | 0 um<sup>3</sup> | |
| $r_{NS}$ | 0.33 hour<sup>-1</sup> | 0.33 hour<sup>-1</sup> | 0.35 hour<sup>-1</sup> | 0.013 hour<sup>-1</sup> | |
| $V_{CS}$ | 976 um<sup>3</sup> | 488 um<sup>3</sup> | 0 um<sup>3</sup> | 0 um<sup>3</sup> | |
| $r_{CS}$ | 0.27 hour<sup>-1</sup> | 0.33 hour<sup>-1</sup> | 1 hour<sup>-1</sup> | 0.0032 hour<sup>-1</sup> | |

#### Cell mechanics

Cells in Gell are mechanically modeled as elastic balls with varied volumes and center positions. Cells adhere to each other while attached and push against each other upon compression. The motion of cell i at position x_i_(t), with velocity v_i_(t), and with a set N_i_(t) of nearby cells can be modeled as (15):

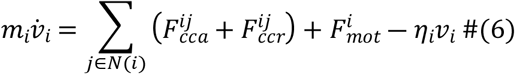

Where F_cca^ij^_ denotes the adhesive force from cell j to cell i, and F_ccr^ij^_ represents the repulsive force from cell j to cell i. F_mot^i^_ accounts for the force related to cell migration. η_i_v_i_ represents the resistance contributed by the local microenvironment, such as fluid resistance and cell-matrix adhesion forces. η_i_ is like fluid drag coefficient and v_i_ is the cell velocity.

The force equilibrates at relatively short time scales relative to the time scale of cell volume change and multicellular patterning. Therefore, we can safely apply the zero acceleration inertialess condition to Eq (6) and explicitly solve v_i_ by:

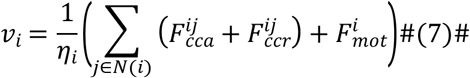

The adhesive force and repulsive force experienced by cell i are modeled as:

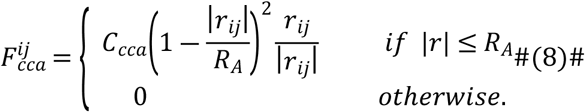

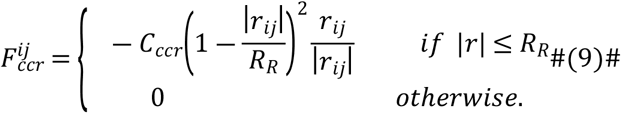

R_R_ is the maximum repulsive interaction distance that equals the sum of the radius of cell i and j. R_A_ is the maximum adhesive interaction distance, which is slightly larger than R_R_ due to the deformability of the two cells. C_cca_ is the cell-cell adhesion parameter and C_ccr_ is the cell-cell repulsion parameter. r_ij_ is a vector pointing from the center of cell i to the center of cell j.

Once the sum of the experienced force is calculated for all cells, the velocity of any cell can be directly calculated. Cell position is then updated using the second-order Adam-Bashforth method:

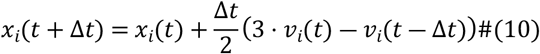

### Extracellular microenvironment

The tumor is surrounded by a complex ecosystem named tumor microenvironment (TME), composed of tumor cells, stromal cells, and other extracellular physical and chemical factors. The mutual and dynamic crosstalk between the tumor and tumor microenvironment, together with the genetic/epigenetic change in tumor cells, are two factors that influence the formation and progression of the tumor(23). In our model, cells can absorb environmental nutrients and release biochemical factors into the extracellular fluid. In addition, critical environmental factors can also regulate cell behaviors. To simulate the spatio-temporal variation of environmental factors during tumor development, we consider a continuous extracellular fluid space and use PDEs to describe the secretion, diffusion, uptake, and decay of diffusive substances such as oxygen and vascular endothelial growth factor. The continuum environment and the discrete cells are explicitly linked. Cell phases, sizes, and positions are treated as static while updating the continuous molecular space and vice versa. The equation for any diffusive substance in the extracellular fluid domain Ω can be written as (24)

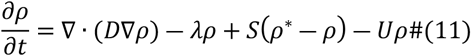

Depending on the problem, the domain boundary ∂Ω can be either Dirichlet or Neumann type. ρ is the substance concentration, ρ* is the saturation concentration, D is the diffusion coefficient, λ is the decay rate, S is the supply rate, U is the uptake rate.

In the provided code, the oxygen concentration is the only considered environmental factor and discrete cells are the only contributor of oxygen consumption. Cells absorb oxygen from the extracellular fluid and regulate their behavior according to local oxygen concentration. The oxygen consumption of each cell is modeled as proportional to both the oxygen concentration and the cell volume. With a cartesian grid, for each isotropic voxel i, the total uptake rate of oxygen equals the sum of the local cell consumption:

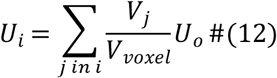

The uptake rate here represents the oxygen concentration decrease rate as a proportion of current concentration, U_o_ is the default oxygen consumption rate of living cells that equals 10 per min. V_voxel_ is the volume of the given voxel, j is the index of cells inside the voxel, and V_j_ is the corresponding cell volume.

In the simulation, each voxel’s total oxygen consumption rate is first calculated according to the position, size, and phase of all the cells. Then the molecular space is updated using the static consumption rate map. With the calculated molecular concentration, cells in the discrete model could read the local oxygen concentration and carry on their stochastic phase transitions according to these values.

For the numerical processing of the PDE, following BioFVM(24), a first-order splitting method is first applied to split the righthand side into simpler operators: a supply and uptake operator and a diffusion-decay operator.

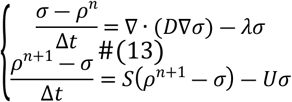

The supply and uptake operators are handled analytically. The three-dimensional diffusion-decay operator is further split into a series of related one-dimensional PDEs using the locally-one dimensional (LOD) method.

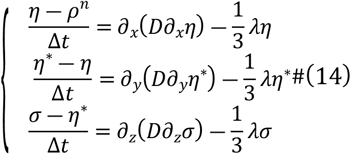

Discretized using the finite volume method, the updated concentration of each strip of voxels for each direction can be obtained by solving Eq (15) using the Thomas algorithm.

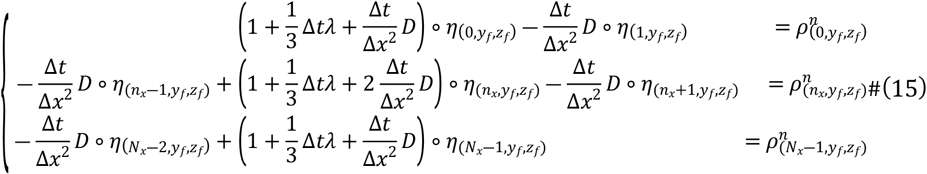

### Implementation Details

Gell is developed using C++ and CUDA. Once initialized on the CPU, all the cellular and extracellular environmental data are transferred to GPU memory. From this point, all the following computation steps are exclusively handled by GPU to eliminate costly back-and-forth data transfer between GPU and CPU. All calculations are on a personal computer (Intel® Core™ i7-7700K and NVIDIA® GeForce® RTX 2080Ti GPU).

#### Cell cycle update

Cell growth, phase transition, and nutrient consumption only involve the current cell status and the environment voxel where the cell locates. These processes are highly parallelizable and can be easily distributed to different threads and be computed by GPU with high efficiency. For processes requiring memory management of structured data, such as cell proliferation and death, fast simulations can also be achieved with only negligible additional computation and memory cost for thread competition avoidance. The overall computational complexity of the aforementioned processes is approximately O(N).

#### Cell mechanics update

N-body interaction simulations can be extremely expensive due to its O(N_cell_^2^) computational complexity. For a large multicellular system with millions of cells, it is computationally impractical just to loop over all the cell pairs, even with GPU(18)(20). Fortunately, the cell-cell mechanical interactions are short-range interactions, making it reasonable to calculate only the forces between neighboring cells, thus reducing the computational complexity to O(Ncell). PhysiCell utilizes the cell-cell interaction data structure (IDS) method(15). A large number of lists are created and maintained to record the indices of cells inside each voxel. The force calculation for each cell only has to loop over the cells inside its nearest 27 voxels, according to the lists. GPU BioDynamo, with its uniform grid method(21), improves memory usage efficiency by replacing the cell index list with the linked list. However, maintaining such cell lists or linked lists is not GPU friendly. GPU does not allow us to dynamically allocate memory in the thread, making it challenging to create lists with dynamic lengths for all mechanical voxels in the IDS method. Suppose GPU memory for cell lists are all preallocated according to the maximum cell density. In that case, memory usage can be highly inefficient due to the high cell density necrotic region. The challenge lies in the maintenance of the linked list for the uniform grid method on GPU. Thread locks are required to update the linked list correctly, but the wrapping mechanism of CUDA could easily create deadlocks during such processes and pause the program indefinitely.

To realize an efficient cell-cell mechanics computation on GPU, we have developed our Voxel-Sorting Method. Cells are stored in array-of-structures, and a Morton code is generated for each cell according to the i, j, k index of its containing mechanical voxel as a key for sorting. Then a fast GPU-based radix sort algorithm(25) of complexity O(N_cell_) is used to rearrange the cell array according to the ascending Morton code value order. After sorting, cells in the same voxel are stored contiguously in the GPU memory, and cells in adjacent voxels are stored relatively close. Such a contiguous memory layout increases the memory fetch efficiency, especially when groups of cells in the same voxel have to be accessed together by the same thread. The array index range for cells inside each voxel is easily determined with O(N_voxel_).

**Fig 1.**
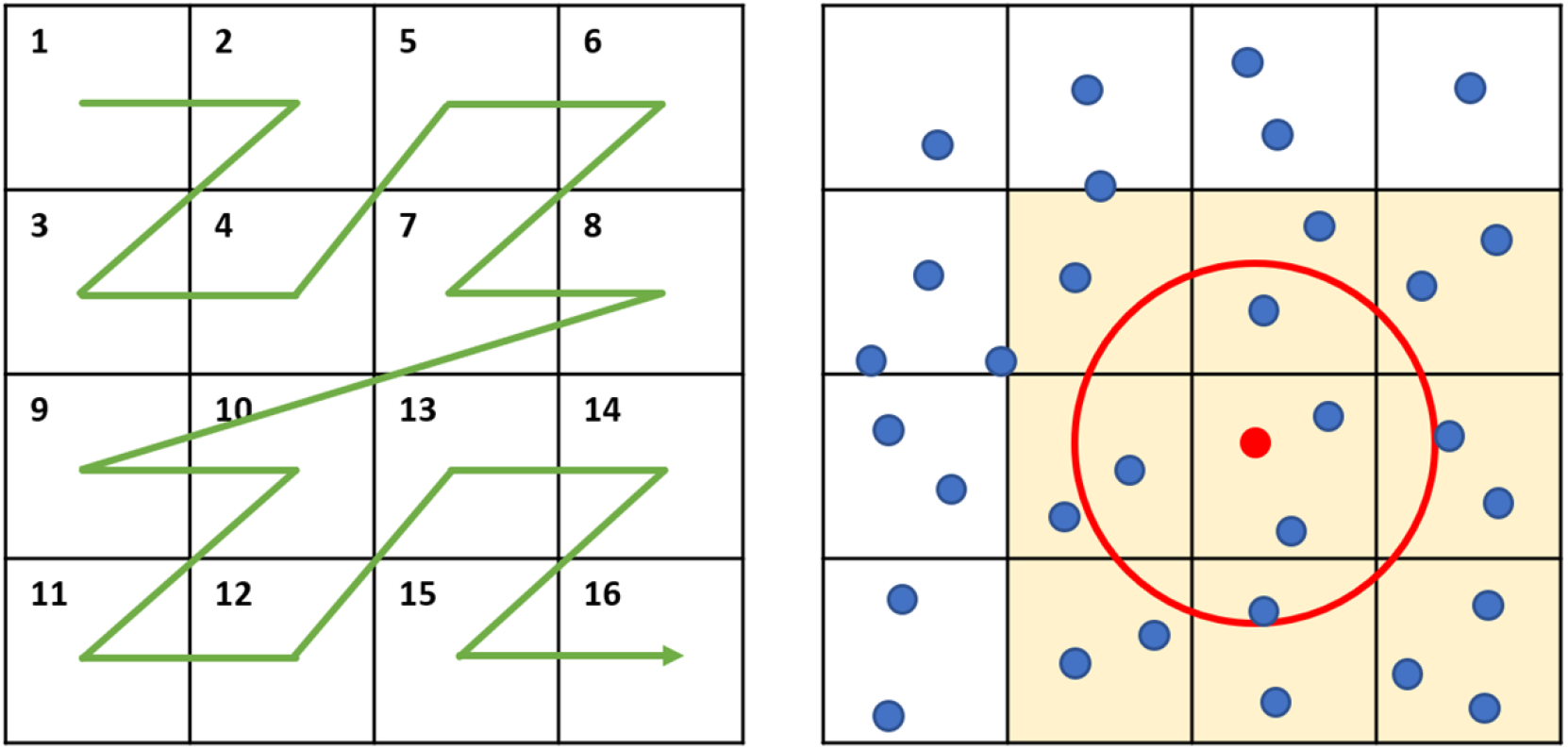
2D illustrations of voxel sorting method. Left: The Morton code maps 2D position to 1D number while preserving the locality of the data points. The Morton code is generated by interleaving the binary x and y pixel index values. Adapted from(26). Right: The force calculation example of a single cell(red). The red circle represents the maximum range of cell-cell mechanical interaction whose radius is shorter than the side length of voxels. The Force module only needs to loop over the cells in yellow voxels to calculate the aggregated force, and the indexes of cells inside these voxels can be easily figured out after the sorting.

As long as the voxel side length is longer than the maximum cell-cell interaction distance, each cell’s aggregated cell-cell mechanical interaction can be calculated efficiently by looping only over cells inside its nearest 27 voxels. Our voxel sorting method achieved a force calculation time complexity of O(N_cell_) and high GPU memory utilization efficiency. An additional advantage of our voxel sorting method is memory access efficiency. With the cells in the same voxel stored next to each other in the GPU memory after voxel sorting, the following 27 times of memory fetch of these cell data can become much more efficient than fully random memory access. After the force calculation, the cell position update can be directly parallelized using the aggregated force information.

#### Extracellular microenvironment update

The problem of spatio-temporal variation of diffusive substances in the extracellular fluid is handled by the LOD solver entirely on GPU. The 3D domain is discretized into N^3^ cartesian voxels of the same size as mechanical voxels to match the cellular level simulation accuracy. At each diffusion-reaction step, the LOD solver converts the concentration update problem along the three axes into N^2^ tridiagonal linear systems, each with N unknowns. Each paralleled thread solves one of these tridiagonal linear systems. The overall computational complexity of each update step is O(N_voxel_).

#### Time scale considerations

Different biological processes evolve at different time scales: the temporal scale of cell colony biology and mechanics is on the order of minutes, while the equilibrium of transport diffusion is achieved in seconds(15). Because the extracellular environment updates substantially faster than cellular evolution, it is computationally inefficient to synchronize the cell simulator and extracellular environment simulator updates. Instead, we first fixed cellular properties while solving the PDEs for the extracellular environment at a higher frequency and then evolved cell phenotype and mechanical interaction at a lower frequency to reduce the simulation cost.

## Results

### Hanging drop spheroid (HDS)

To compare the computational performance of our simulation framework with existing simulation software, we simulated a benchmark problem of hanging drop spheroid growth(15). In the HDS simulation, a suspended multicellular aggregate is cultured in the middle of a growth medium with oxygen supplied through diffusion from the domain boundary. All the Gell simulations and PhysiCell simulations share identical simulation settings. The simulation started with 2347 cells and evolved to one million cells after 19 days of cultivation. The simulation domain contains one million isotropic voxels with a side length of 25 um. Cell mechanics and phase update is calculated every 0.1 minutes, and the diffusion-reaction of oxygen in the extracellular fluid is solved every 0.01 minutes.

The evolution of cell number and spheroid radius over time is shown in Fig 2. The result shows no necrosis development until the spheroid reaches a certain radius when oxygen diffusion from the outer rim becomes insufficient to support the inner cells. The radius growth curve exhibits a linear shape due to the approximately constant viable rim thickness, which agrees with the simulation result and theoretical prediction of PhysiCell(15).

**Fig 2.**
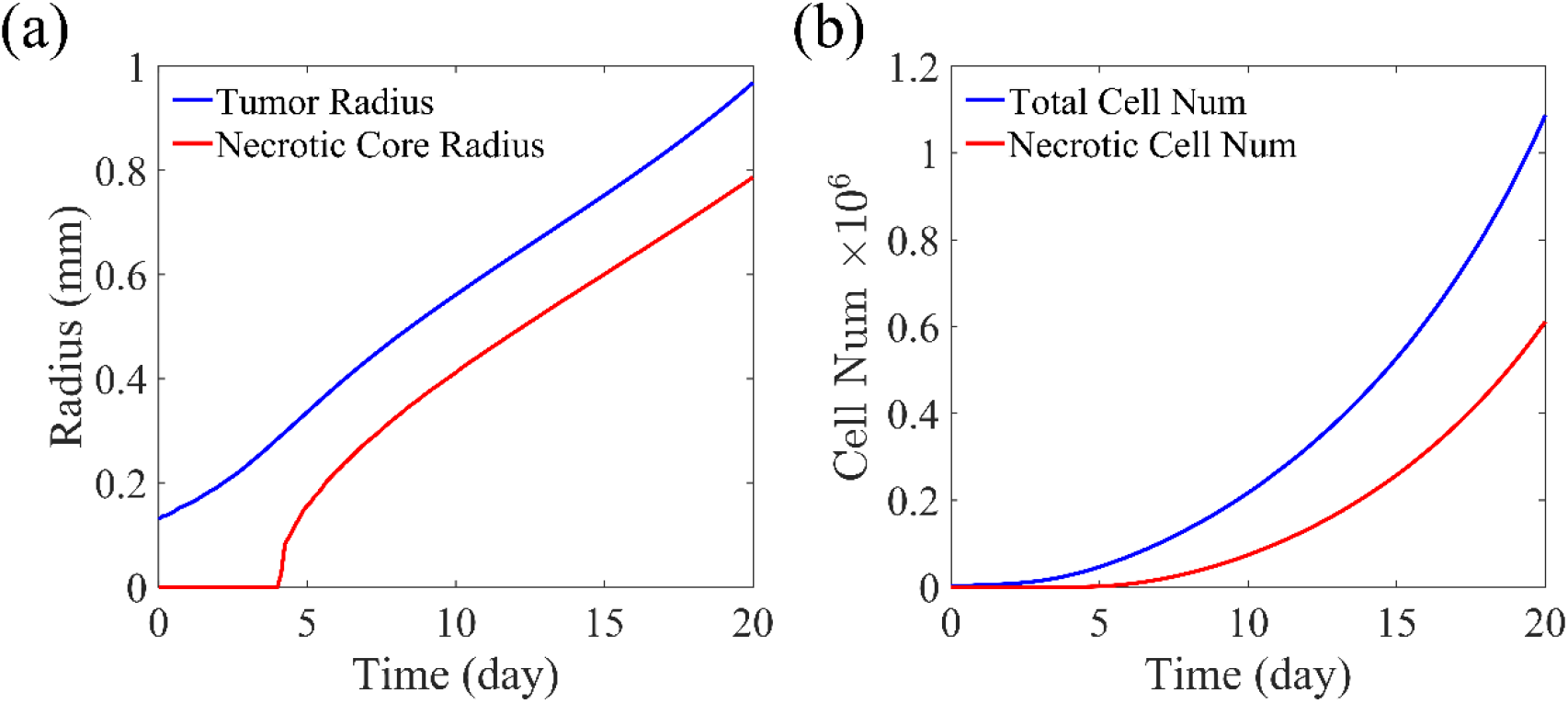
Simulation result analysis of Gell. (a) Whole tumor radius and necrotic core radius change over the HDS growth. (b) Number change of total tumor cells and necrotic tumor cells over the HDS growth.

The rendered images of the simulation results are shown in Fig 3. There is a clear viable rim of actively proliferating cells at the outer shell of the spheroid and a necrotic core in the center (Fig 3.a). Additionally, the subtle cell-cell mechanical adhesion, crack like microstructures successfully emerged in the necrotic core enter. The simulation results of Gell agree with PhysiCell and in vitro experiments(15).

**Fig 3.**
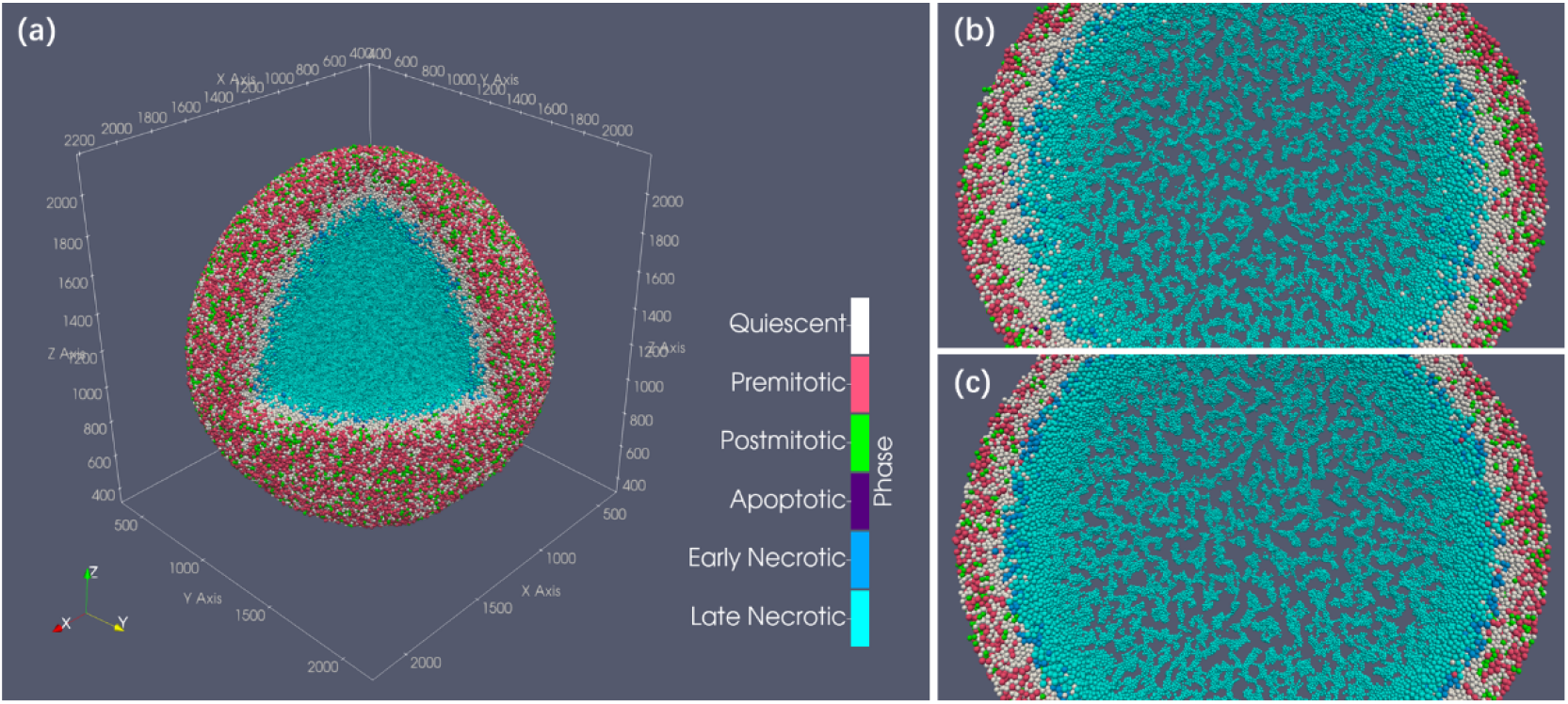
HDS simulation result after 450 hours. (a) Cell cluster generated by Gell. (b) 60 um thick central slice of the HDS simulation result shows the microstructure of the necrotic core of Gell simulation. (c) central slice of PhysiCell showing identical microstructure. Both spheroids have a radius of 1.87 mm.

Gell completed the entire simulation process in 47 minutes using the personal computer. As a comparison, the state-of-the-art CPU-based paralleled simulator PhysiCell used 119 hours for the same simulation on the same personal computer. In other words, Gell is 150X faster for the cell simulation problem of this scale.

Benefiting from the accelerated computation, we were able to explore the parameter space without agonizing pain. We surprisingly found that the crack pattern of the necrotic core has little to do with the two-stage necrosis process of tumor cells. The central slice of the spheroid of non-swelling tumor cells shows almost identical microstructure patterns (Fig 4.b). However, this system ends up with a slightly smaller size, fewer cells, and a higher overall necrotic debris density. This suggests that the swelling process of tumor cells could facilitate spheroid growth by pushing the viable cells towards the more oxygenated outer regions. Fig 4.c shows the spheroid of tumor cells with the cell-cell adhesion suppressed. The cell-cell adhesion factor Ccca is decreased to one-fourth of the reference value. The weak adhesion discourages the gathering of necrotic cells leading to a more scattered distribution of smaller necrotic cell clusters with more minor interleaving cracks. Fig 4.d is a spheroid with the cell-cell adhesion enhanced by quadrupling Ccca. The strong adhesive force helps form the massive necrotic debris clusters and promotes a significantly higher cell density that intensifies the oxygen competition between tumor cells and ultimately hinders tumor growth. Such pattern differences in necrotic core microstructures that emerged in the simulation are also observed in in vitro experiments (27), as shown in Fig 5. Two spheroids of the same melanoma cell line (A2508) form clustered (Fig 5.a) and scattered (Fig 5.b) necrotic cores, respectively. The exact differences in cell treatment and mechanisms of pattern formation are not described in the original literature. However, our simulations hint that cell-cell adhesion could be an important factor that dramatically affects the spheroid morphology, especially the necrotic core microstructure.

**Fig 4.**
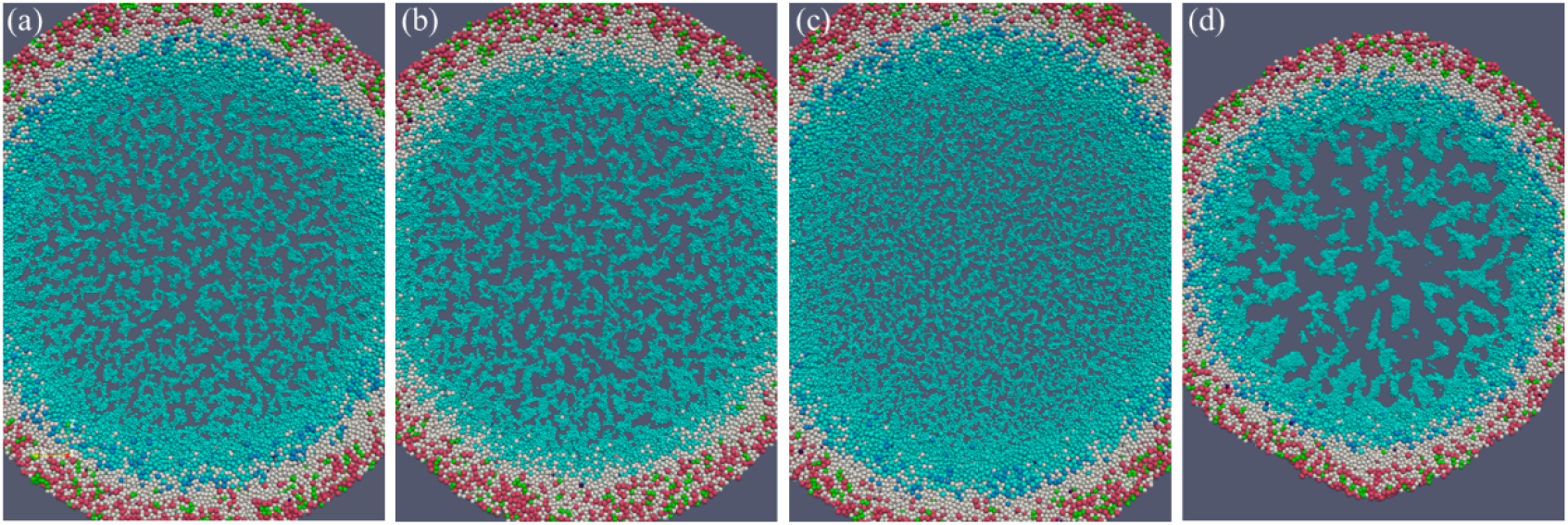
HDS simulation with altered phenotype. The 60-um thick central slices of simulated spheroids with various cellular mechanical properties. All the spheroids start with a small cluster of 2347 randomly placed cells, and the cultivation duration is 450 hours. (a) The reference spheroid ends up with 0.9 million cells and a diameter of 1.87 mm. (b) Spheroid of tumor cells with no swelling during early necrosis, with 0.9 million cells and a diameter of 1.8 mm. (c) Spheroid of tumor cells with the cell-cell adhesion suppressed, with 1.0 million cells and a diameter of 1.97 mm. (d) Spheroid of tumor cells with the cell-cell adhesion enhanced, with 0.66 million cells and a diameter of 1.53 mm.

**Fig 5.**
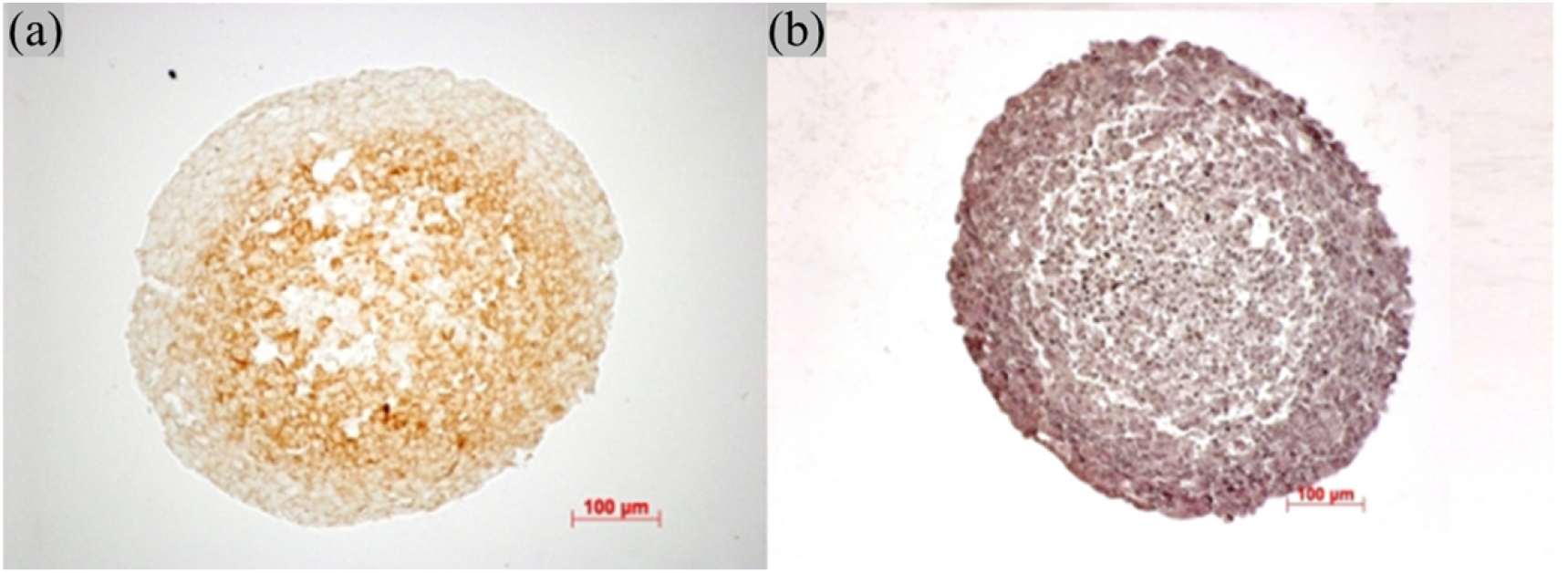
Melanoma cell line spheroids. Two spheroids of the same melanoma cell line (A2508) show distinct pattern differences in necrotic core microstructures due to differences in cell treatment. Images are adapted from (27), and treatment details are not mentioned in the original literature. (a) A pimonidazole stained spheroid. (b) A hematoxylin and eosin stained spheroid. Adapted from (27) with permission.

Simulations have the potential to depict a causal and mechanistic path from certain microscopic cell properties to the development of qualitative macroscopic morphology and provide insights into real-world phenomena. However, it can be very time-consuming to explore the parameter space and to improve the models iteratively. Gell could help these studies by dramatically increasing the computational speed.

Besides the baseline HDS simulation task, further comparisons of simulation performance with varying initialized cell numbers (Fig 6.a) and domain sizes (Fig 6.b) suggest that Gell is consistently around two orders of magnitude faster than multi-thread PhysiCell on the personal computer. For serial PhysiCell using only one thread, Gell can be almost 400× faster (Fig 6.c). The default simulation setting for the performance comparison contains one million 25×25×25 um isotropic voxels and one million living cells. Cells are randomly initialized in a sphere with a specific cell density. The acceleration ratios of these tests differ from the whole HDS simulation because the HDS simulation has a more heterogeneous cell distribution and is closer to the equilibrium states after prolonged mechanical interactions.

**Fig 6.**
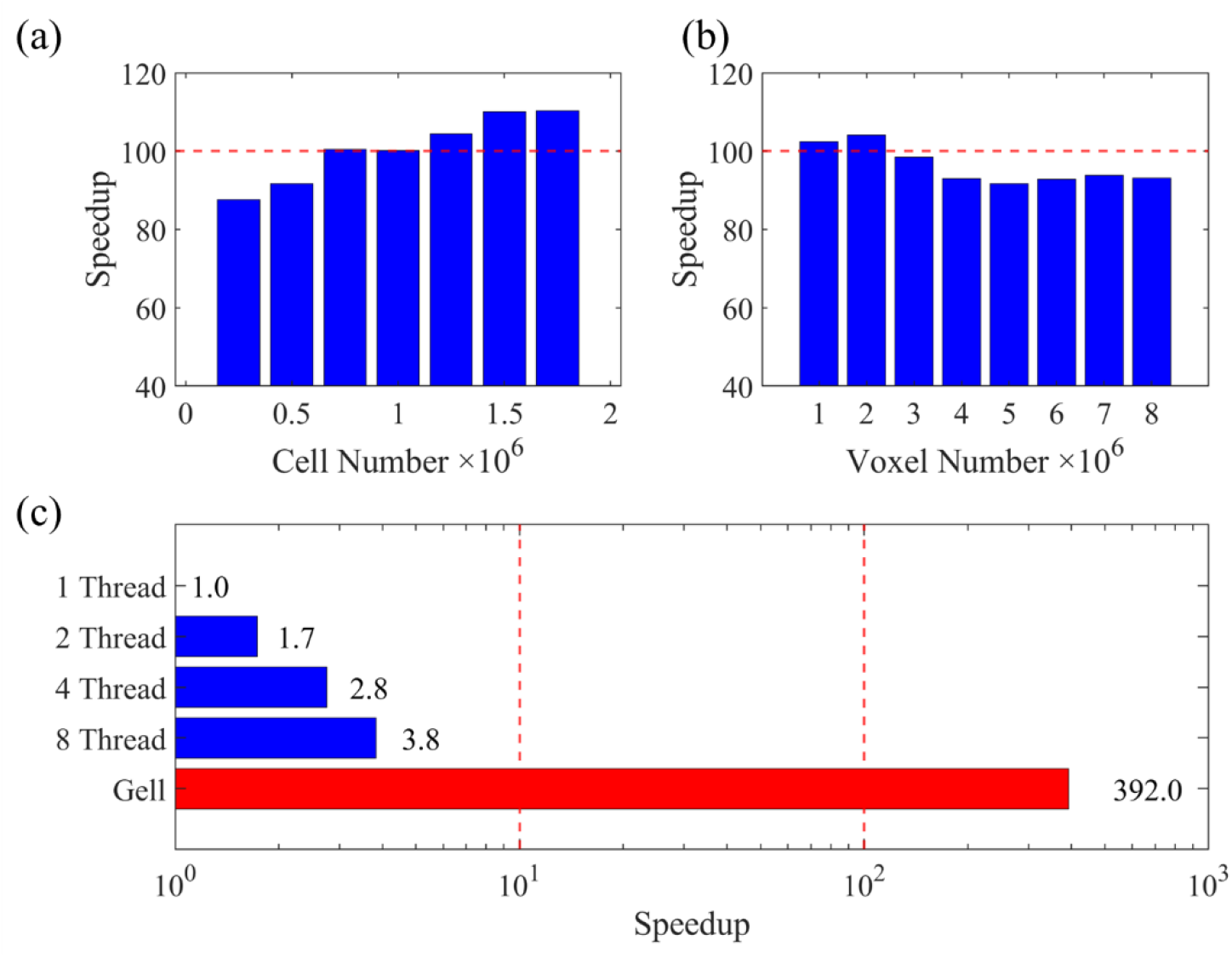
Gell simulation speedup with respect to PhysiCell. Gell simulation speedup with respect to PhysiCell with varied cell number (a), domain size (b), and PhysiCell CPU thread numbers (c).

Additionally, Gell can complete the simulation with its maximum memory footprint being only one-tenth of that of PhysiCell’s without sacrificing accuracy and system complexity (Table 4).

**Table 4.** Memory footprint comparison.

|  | PhysiCell | Gell |
| --- | --- | --- |
| Memory Footprint of HDS Simulation | 6360 | 500 |
| Memory Footprint per Additional Million Cells (MB) | 5384.2 | 118.0 |
| Memory Footprint per Additional Million Voxels (MB) | 742.8 | 15.38 |

Memory footprint comparison of Gell and PhysiCell per additional million cells and per additional million domain voxels in the HDS simulation task. Our model is able to simulate the problem with a much lower memory occupation.

### Ductal carcinima in situ (DCIS)

Ductal carcinoma in situ is non-invasive breast cancer that grows within the lumens of the mammary duct(28). DCIS itself is not hard to treat, but as a precursor to invasive ductal carcinoma with a high incidence rate (26.6 per 100000 women(29)), its development and progression raise the interest of many modelers(30) (31) (32) (33) (34). Following the work of Paul Macklin (30), we created a hybrid DCIS simulation example to further illustrate Gell with a more complex task. A cluster of tumor cells is placed at the dead-end of a fixed single-opening duct. The movement of tumor cells is confined within the tube lumen, and the oxygen is supplied through diffusion across the duct wall into the lumen. Breast ducts have a typical radius ranging from 100 um to 200 um (32); therefore, we simulated three growth scenarios of ductal carcinomas in situ with the respective duct radius of 100 um, 150 um, and 200 um.

**Fig 7.**
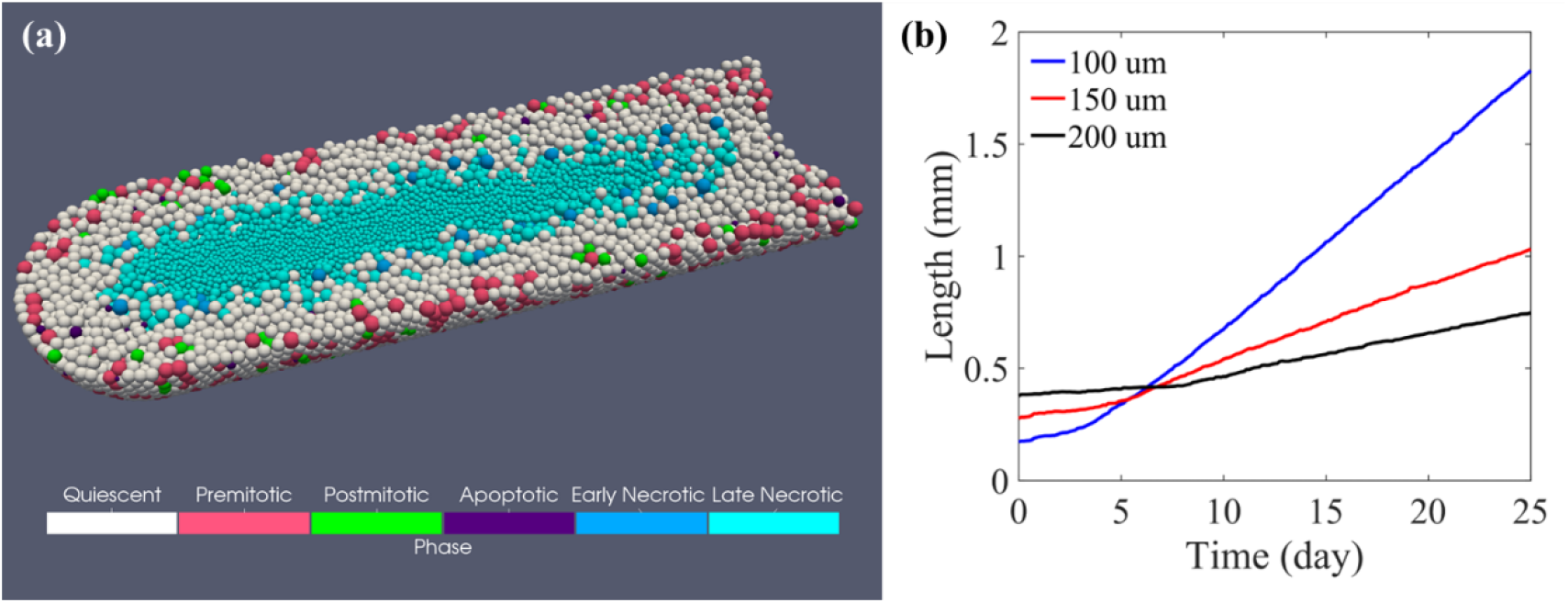
Simulation results of DCIS development. (a) Ductal carcinoma in situ simulation with duct radius of 150 um. (b) Linear DCIS growth under various duct radius conditions.

With the same cell model as in the HDS simulation and a Dirichlet boundary condition of 7.2 mmHg oxygen concentration applied to the duct surface, the average rate of DCIS advance for ducts of radius 100, 150, and 200 um is 77.0, 32.7, 19.0 um/day, respectively. Our simulation results show a linear growth speed of DCIS, and the tumor advance rate has an inverse relationship with the duct radius, which agrees with the clinical observations and other computational studies in 2D(30) or 3D(35).

### Performance testing

#### Individual module time cost

For performance assessment, we first evaluated our simulator with a randomly initiated spherical cell cluster with one million cells. The simulation domain contains one million 25 um-long isotropic voxels. The time cost of each module is listed in Tab. 4. With the entire calculation paralleled on GPU, Gell maximizes computational efficiency in all the simulation modules. Meanwhile, unnecessary data transfer between CPU and GPU during the simulation is eliminated, ensuring a low delay between modules.

In the table, the cell-sorting-related process appears to be slow, but in practice, its impact on the overall computation is small. Firstly, the cell sorting module is less invocated than many other modules. The simulation faces a multiscale problem. The cell phase is updated every few simulation minutes, cell motion is updated every few seconds, and diffusion-reaction of oxygen is updated more than once per second. In this case, the simulation is approximately equally dominated by the cell-motion- related modules and the diffusion-reaction model. Secondly, the sorting process accelerates the rest of the calculations. Take the oxygen consumption model as an example. This kernel is a part of the reaction-diffusion module that calculates each cell’s oxygen consumption rate and adds the value to the total consumption rate of each corresponding voxel. This kernel works with both unsorted and sorted cell structures. Changing from unsorted random memory access to sorted contiguous memory access, this module’s time cost per invocation is reduced from 0.345 ms to 0.170 ms by 51%. The force calculation module is expected to benefit most from the high data fetch efficiency of the voxel sorting method. Because force calculation does not work with unsorted data structures, the speed comparison cannot be performed. Nevertheless, the longer time spent on cell sorting benefits the overall computation and is a worthy investment.

#### Performance scaling

Unaffordable memory occupation is another potential barrier for large-scale cell-based simulations. Memory efficiency is another design goal to make Gell a suitable software for large-scale simulations on widely available devices. The GPU memory footprint of the HDS simulation with one million voxels and one million cells is limited to 500 MB, and the memory occupation increase with the cell number and domain size is linear and slow, as shown in Table 5. This enables Gell to fit extremely large-scale problems into a modern personal computer easily.

**Table 5.** The execution time cost of each simulation module in Gell.

| Module | Time cost per invocation |
| --- | --- |
| Morton code calculation | 0.303 ms |
| Cell sorting | 6.041 ms |
| Force calculation | 3.613 ms |
| Cell movement | 0.360 ms |
| Reaction-diffusion update | 0.542 ms |
| Phase update with birth and death | 4.744 ms |
| Memory copy between CPU and GPU | 0.011 ns |
| Others | 0.387 ms |

Values are averaged over 1000 simulation steps with the time step for phase update, mechanics update, and diffusion-reaction update all set to be the same. Short time such as the time cost of memory copy during a simulation, is estimated by comparing a 1000-step simulation with another 2000-step

We tested Gell’s ability to handle ultra-large-scale problems with a hypothetical ultra-large spheroid model. In reality, the size of hanging droplets is diffusion constrained. Larger in vivo tumors inevitably involve angiogenesis and supporting tumor vasculature. However, angiogenesis and oxygen transportation simulations are beyond the scope of the current work. Instead, we simulate a series of hypothetical huge hanging droplet development problems in a huge domain with varying initial cell numbers, each for one hour. As shown in Fig. 8, Gell has a linear computational cost scaling with the cell number. Simultaneously, the Gell GPU memory footprint peaked at 4392 MB, showing high efficiency in memory and computational source usage. This linear time complexity and low memory occupation demonstrate that Gell can handle potential ultra-large-scale problems.

**Fig 8.**
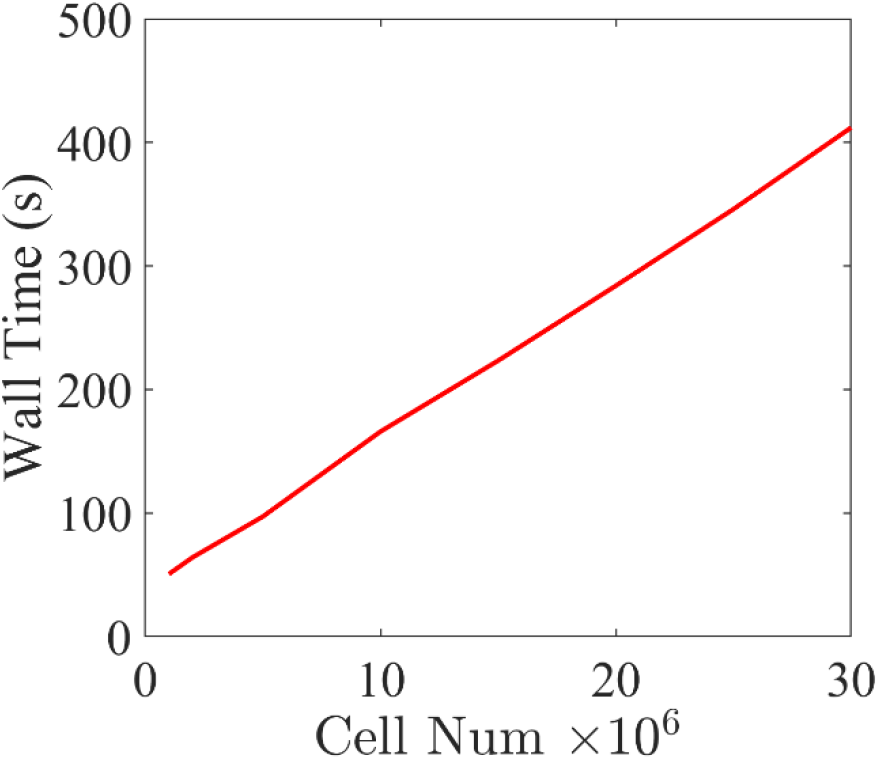
Performance scaling of Gell. Time cost for one hour’s mechano-biological process simulation with varying cell number. The domain contains 250 ×250 ×250 25 um-long isotropic voxels.

## Discussion

Cell simulation is driven by the need to model larger and more complex digital tumors parallel to human tumors containing billions of cells. Simulation of such complex systems has been limited by the modeling accuracy of the biology and computational capacities. Existing tools such as Biocellion(16) for such large-scale simulations are close-sourced and require expensive CPU clusters. Open source tools, including Physicell (15) and BiodynaMo (17) require high-end CPU clusters to perform computation in practical time. In theory, GPU is well suited to manage the large parallel components of cell simulation. However, due to the differences between GPU and CPU architecture, a direct translation of a CPU-based cell simulation code to a GPU-based one is inefficient. Specifically, the global cell-cell interactions have a time complexity of O(N^2^), severely limiting the number of cells that can be modeled. To alleviate the challenge, a neighboring cell list needs to be maintained, which is difficult for GPU memory due to its dynamic nature. Other than this drawback, the incomplete translation of computation from CPU to GPU requiring frequent data transfer between them is another common limitation. As a result, the threshold of performing cell simulation to a size that is relevant to the small tissue scale remains out of reach for many biological researchers.

To overcome the challenges, we employed several novel techniques to more thoroughly exploit the unique computational architecture of GPU and improve the simulation performance while minimizing memory usage.

As the computational cost of cell-cell interaction rises quadratically with the cell number, we developed a GPU-friendly voxel sorting method that efficiently handles the short-range cell-cell interaction modeling and improves the whole simulation’s memory access efficiency. We also implemented a fully GPU-based LOD solver for the spatio-temporal variation of diffusive substance distribution in extracellular water. As a result, we optimized the evolution algorithm to achieve linear computational complexity O(N) while minimizing the memory footprint. In the numerical implementation, we fully exploited the parallel architecture of modern GPU and different types of GPU memory for high computational speed and low memory access overhead. The computation is nearly 100% on GPU, avoiding slow data transfer between CPU and GPU memories.

Our GPU implementation significantly outperformed CPU methods and led to almost 400X speedup over the single thread version of the well-established CPU simulator on a personal computer. The acceleration is the highest among existing GPU-powered simulators. The acceleration, in combination with the low memory footprint, makes Gell easily accessible to biology researchers. The easy-to-access platform would facilitate the fast prototyping of hybrid models and hypothesis testing for large-scale problems.

As a future research direction, Gell can be scaled to multiple GPUs for larger problems on the order of 10^9^ cells. As previously eluded to, simulation of tumor vasculature and other scaffolding cells such as the stromal cells and immune cells would be necessary for a biologically relevant model. Besides our effort to incorporate these extremely complex biological processes for digital tumor twins, we support our peers to join the effort by providing and updating Gell as a user-friendly open-source tool. The source code can be found at https://github.com/PhantomOtter/Gell.

## Notes

### Competing Interest Statement

The authors have declared no competing interest.

